# Lost in translation: No effect of repeated optogenetic cortico-striatal stimulation on compulsivity in rats

**DOI:** 10.1101/2020.10.19.345942

**Authors:** Amanda R. de Oliveira, Adriano E. Reimer, Gregory J. Simandl, Sumedh S. Nagrale, Alik S. Widge

**Author notes:** These authors contributed equally to this work (Amanda R. de Oliveira, Adriano E. Reimer). Corresponding Author: Alik S. Widge, Department of Psychiatry, University of Minnesota, Minneapolis, MN 55455.

## Abstract

**BACKGROUND:** The orbitofrontal cortex-ventromedial striatum (OFC-VMS) circuitry is widely believed to drive compulsive behavior. Hyperactivating this pathway in inbred mice produces excessive and persistent self-grooming, which has been considered a model for human compulsivity. We aimed to replicate these findings in outbred rats, where there are few reliable compulsivity models.

**METHODS:** 27 male Long-Evans rats implanted with optical-fibers into VMS and with opsins delivered into OFC received optical stimulation at parameters that produce OFC-VMS plasticity and compulsive grooming in mice. We then evaluated rats for compulsive self-grooming at six timepoints: before, during, immediately after and one hour after each stimulation, one and two weeks after the ending of a 6-day stimulation protocol. To further test for effects of OFC-VMS hyperstimulation, we ran animals in three standard compulsivity assays: marble burying, nestlet shredding, and operant attentional set-shifting.

**RESULTS:** OFC-VMS stimulation did not increase self-grooming or induce significant changes in nestlet shredding, marble burying, or set-shifting in rats. Follow-on evoked potential studies verified that the mouse protocol did alter OFC-VMS synaptic weighting.

**CONCLUSIONS:** In sum, although physiological changes were observed in the OFC-VMS circuitry, we could not reproduce in a strongly powered study in rats a model of compulsive behavior previously reported in mice. If optogenetic effects on behavior do not reliably transfer between rodent species, this may have important implications for designing rodent-to-human translational pipelines.

## Introduction

Compulsions – maladaptive patterns of repetitive, inflexible cognition and behavior – are a key feature of numerous mental health conditions, including obsessive-compulsive disorder (OCD), trichotillomania, skin picking disorder, eating disorders, addiction, anxiety, and depression (1–4). Mental disorders associated with compulsive behaviors cause significant distress and, because they include various forms and degrees of manifestation, they still are difficult to diagnose and treat (3,5–7). Thus, better understanding the pathophysiology of compulsive behavior could optimize therapeutic approaches for multiple disorders.

Compulsivity is increasingly understood as resulting from failures of information flow in the neural circuits that govern self-regulation, motivation and learning (2,8–10). Clinical and preclinical studies suggest that compulsivity is governed by the frontal lobes in a complex interaction with subcortical systems that include midline thalamic nuclei, striatal regions, and the mesocorticolimbic dopamine system (11–14). Consequently, abnormalities in the function of this cortico-striatal circuitry can contribute to compulsive/inflexible behavior, especially in complex and changing environments. For instance, accumulating evidence points to dysregulation of the cortico-striato-thalamo-cortical (CTSC) circuitry as a cause of OCD, one of the most common diseases of compulsive behavior (10,15–17). Orbitofrontal cortex-ventromedial striatum (OFC-VMS) hyperconnectivity is particularly thought to drive compulsivity (10,18–20). However, it is uncertain whether OFC-VMS dysfunction is sufficient to produce compulsivity on its own.

Non-human animal studies can contribute to the identification of relevant circuits such as the OFC-VMS pathway. The main barrier to these studies, however, has been the lack of a clear link between human disease and animal behavior models, particularly in those models that respond to manipulations of the CTSC circuits. Self-grooming has been proposed as an animal model of inappropriate compulsive behavior (21,22). Grooming is a ritualized sequence of coordinated movements that include licking, scratching and wiping, aimed at cleaning fur and skin (23–25). Self-grooming has clear adaptive functions in certain situations (26,27), but it is displaced and mal-adaptive in others. It may appear, for example, when the animal is placed in atypical or stressful situations, such as when exposed to the elevated plus-maze test (26,28).

In mice, hypergrooming is specifically linked to the OFC-VMS circuit. Ahmari et al. (20) used an optogenetic strategy to hyperactivate the OFC-VMS circuit. Repeated optogenetic stimulation of OFC-VMS projections in inbred mice produced excessive selfgrooming behavior. This grooming was specifically the result of OFC-VMS plasticity: it appeared only after multiple days of stimulation, persisted in the absence of stimulation, and was reversed with chronic administration of fluoxetine, a drug considered a first line pharmacotherapy agent for OCD. The change was specific to compulsivity: the model showed no differences from unstimulated mice in sensorimotor filtering (pre-pulse inhibition) or anxiety-related activity (elevated plus-maze and open-field tests).

A major open question is whether the link between induced OFC-VMS hyperconnectivity and compulsivity can translate across species. Most of the literature on induced compulsivity is in mice, given the large genetic toolbox e.g., the HoxB8 (29), Slitrk5 (30) and SAPAP3 mutants (31). Mice, however, are not as able to participate in complex cognitive tasks, compared to larger rodents such as rats (32,33). Rats are more similar to humans, whether behaviorally □ rats provide a better model system to study alcohol relapse, for example (34), genetically (35), or in central nervous system receptor biology (36). Rats present a larger brain, facilitating electrophysiological and imaging studies while allowing better modeling of clinical interventions such as deep brain stimulation (DBS) (37,38). Finally, if a mouse manipulation does not produce similar behavioral effects in rats, it suggests that the manipulation may not translate well to human behavior either.

We tested whether the OFC-VMS hyperstimulation approach could translate from mice to rats. Since rats might express compulsivity differently compared to mice, and since grooming is only one task-free method for assessing compulsivity, we explored this translation across a range of compulsive behaviors. Other repetitive behaviors considered compulsive-like in rodents include hiding objects and shredding materials recurrently (22,39,40). Thus, we tested the same approach across grooming, marble burying, and nestlet shredding tests, which evaluate repetitive behaviors that may become compulsive (41,42).

An important aspect of human compulsivity that has not been well-modeled in animals is the impairment of behavioral flexibility, meaning deficits in behavior shifting in response to changing environmental contingencies (43–46). Behavioral flexibility often involves withholding a pre-potent (default) response in favor of a more adaptive choice, which in the context of compulsivity could mean avoidance of a previously-emitted behavior. As with the broader construct of compulsivity, flexibility depends on a network that includes OFC, prefrontal cortex and dorsal anterior cingulate cortex (47–50). Flexibility can be tested across species with set-shifting tasks (17), and these tasks have revealed cortico-striatal deficits in patients with OCD (51,52). Similarly, DBS of the ventral capsule/ventral striatum, an advanced surgical therapy for compulsivity, can increase behavioral flexibility (53). Therefore, in addition to the task-free behaviors above, we evaluated the effects of OFC-VMS stimulation on a set-shifting task that tests behavioral flexibility.

Here, we show that in contrast with previous findings in mice, we were not able to induce repetitive behaviors of any type in rats using OFC-VMS repeated stimulation. We also could not disrupt behavioral flexibility in rats by manipulating OFC-VMS circuitry.

Follow-on electrophysiologic studies showed that the lack of behavioral effects could not be explained by an inability of the mouse protocol to alter OFC-VMS synaptic weighting in rats, suggesting that there are fundamental differences in the outcomes of striatal plasticity even between model species.

## Methods and Materials

### Animals

Twenty-seven male Long-Evans rats (250-300 g) were obtained from Charles River Laboratories (Wilmington, MA) and housed in the McGuire Translational Research Facility of the University of Minnesota, Minneapolis, MN, under a 10:14 dark/light cycle (lights on at 07:00 h). We selected our animal counts to be adequately powered to replicate the primary behavioral outcome of the original report of OFC-VMS hyperstimulation (20). Sample size calculations were conducted using GLIMMPSE 2.0.0 (54), with the effect size as observed in Ahmari et al. (20). With the desired power set at 95% and the type I error rate at 5%, to detect a main effect of group, the analysis revealed that a minimum sample size of 16 rats (8 per group) was required (power = 0.956). We targeted above this to account for potential animal exclusions (e.g., due to misplaced injections) and/or a smaller effect size in rats.

Rats were first acclimated for 5-7 days in the animal colony room and, subsequently, were handled for three consecutive days for 5 min/day to familiarize them with the experimenter before the beginning of the procedures. At the end of each of these handling sessions, the rats received five reward pellets (45 mg grain-based pellets; Bioserv, Flemington, NJ) in their home cages to forestall neophobic reactions in the set-shifting operant chambers. Rats had free access to food and water, except during the behavioral training and testing of the set-shifting protocol, when rats were food restricted. For this, animals were restricted to 10 g of standard laboratory rat chow per day until body weight was reduced to 85-90% of the original (after 5 days of food restriction, approximately). At this time food was increased to 15 g per day and their weights maintained at this level without further reduction until completion of the set-shifting sessions. All experiments were approved by the University of Minnesota Institutional Animal Care and Use Committee (protocol number 1806-35990A) and comply with the National Institutes of Health guidelines.

### Surgical procedures

Each rat was deeply anesthetized with 3-4% isoflurane in an induction chamber and was mounted in a stereotaxic frame (maintained on 0.5-2% isoflurane for the duration of the surgery). For opsin delivery, a small opening was made over the left ventromedial orbitofrontal cortex – OFC: VO and MO, as in Ahmari et al. (20) – for the injection of adeno-associated virus (AAV). Injected AAV carried the gene encoding channel-rhodopsin (ChR2) fused to enhanced yellow fluorescent protein (eYFP) under the calcium- and calmodulin-dependent protein kinase II (CaMKII) promoter (AAV5-CaMKIIa-hChR2(H134R)-eYFP, University of North Carolina Viral Vector Core Facility, Chapel Hill, NC). Rats that received virus lacking the ChR2 sequence were used to control for any nonspecific effects of the viral transfection or light stimulation (AAV5-CaMKIIa-eYFP). As in the original study (20), viruses were injected into the left OFC (4.8 mm AP, 0.8 mm ML, 4.5 mm DV) using a 33-gauge Hamilton syringe and a microinjector. Injections (1.0 μl from a 4.1×10^12 vg/ml) occurred over 10 min (0.1 μl/min) followed by an additional 5 min to allow diffusion of viral particles. The use of a CaMKIIa promoter enables transgene expression favoring pyramidal neurons when injected into the neocortex, with a level of transduction efficacy and expression of ChR2 of about 90% (55–58).

For the chronic optical-fiber implant, a small skull opening was made over the left ventromedial striatum (VMS), as in the original study (20). We implanted an optical fiber over the VMS (1.28 mm AP, 2.6 mm ML, 5.8 mm DV) that permitted optical stimulation. Optical fibers were constructed in-house by interfacing a 20 mm piece of 200 μm, 0.66 numerical aperture optical fiber with a 10 mm ceramic ferrule (fiber extending 10 mm beyond the end of the ferrule). Fibers were attached with epoxy resin into ferrules, then cut with a diamond pen and polished. After construction, all fibers were calibrated to determine the percentage of light transmission at the fiber tip that interfaces with the brain; all fibers had > 75% efficiency of light transmission. Several small burr holes were drilled around the perimeter of the exposed skull surface to accept small anchor screws. The fiber was then fixed to the skull and screws with dental cement.

To evaluate the effects of optogenetic stimulation on OFC-VMS plasticity, a second study added recording electrodes. Along with the opsin delivery at OFC and optical-fiber implant at VMS, during stereotaxic surgery, twisted bipolar platinum electrodes with a ground wire (MS333/8-BIU/SPC, Plastics One, Roanoke, VA) were implanted in both regions for stimulation and recording. For the OFC implant, the electrode was lowered 15 min after the end of the AAV injection; for the VMS implant, the electrode was attached to the optical fiber, being set 200 μm below the tip of the fiber. Once the electrodes were placed, ground wires were wrapped to a nearby screw. After surgery, rats were returned to their home cages and allowed to recover for at least 7 days before any behavioral experimentation. A minimum of 5 weeks (for stable viral expression) elapsed before the beginning of the optical stimulation (Fig. 1A-B).

**Fig. 1.**
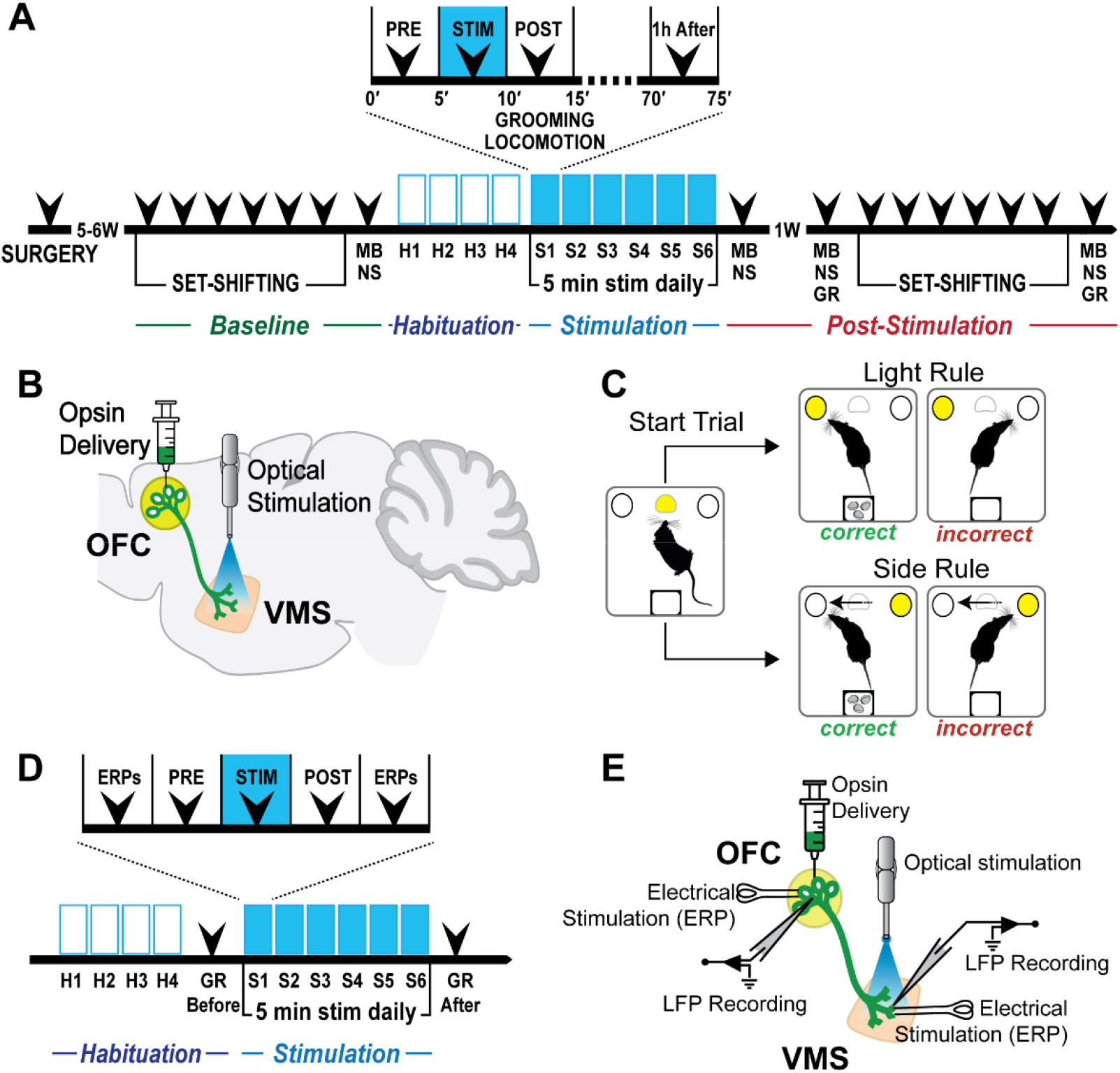
Experimental design for behavioral and electrophysiological evaluation. **A)** Timeline used for repeated stimulation of OFC-VMS projections for the behavioral experiments. H = habituation, S = stimulation, GR = grooming, MB = marble burying. NS = nestlet shredding. **B)** Schematic diagram indicating localization of ChR2-eYFP injections into the OFC and optical fiber implant and stimulation in the VMS. This restricts activation to OFC terminals in VMS. **C)** The operant set-shifting task. Rats had to shift their response patterns between two distinct perceptual discrimination rules: light or side, to receive a reward. Performance according to the light rule required the rats to poke the illuminated nosepoke hole, regardless of its spatial location. Performance according to the side rule required a response at the nosepoke hole at a designated spatial location, regardless of the illumination. **D)** Timeline showing timepoints for evoked potential (ERP) measurement. **E)** Schematic diagram indicating localization of ChR2-eYFP injections into OFC, optical fiber implant in the VMS, and electrical stimulation and recording in both OFC and VMS.

### Optogenetic stimulation

The protocol used for repeated cortico-striatal stimulation followed that described by Ahmari et al. (20). After waiting 5-6 weeks for viral expression, the chronic optical fiber implant was connected to an armored optical patch cable, which in turn was connected to a 1×1 rotary joint optical commutator via FC adaptors and coupled to a 465 nm blue LED. Optical stimulation was controlled by a pulse generator (Pulse Pal, Sanworks, Stony Brook, NY) connected to the LED driver (Plexon, Dallas, TX). For four consecutive days before the beginning of optogenetic stimulation, rats were connected to the patch cord for 15 min per day for habituation (no stimulation was given). The rats” behaviors were recorded by video cameras positioned above and laterally to the cage. On the following 6 days, we repeatedly delivered steady optical stimulation, activating the viral optical payload by bright blue light (5 min/day, 6 consecutive days, 10 ms pulses, 10 Hz, 7 mW), to rats placed in the test chamber. This delivered approximately 55 mW/mm^2^ at the fiber tip, well above the ChR2 opening intensity. Self-grooming behavior was evaluated before, during, immediately after, and one hour after each stimulation, and at one and two weeks after the 6-day stimulation protocol. Marble burying tests, nestlet shredding tests, and operant attentional set-shifting task sessions were performed before and after the 6-day repeated opto-stimulation.

### Self-grooming behavior

Self-grooming behavior was recorded with digital video cameras and scored by a single rater who was blind to the group assignment. Self-grooming was evaluated for 5-min periods: before (pre), during (stim), immediately after (post) and one hour after (1h after) each stimulation. One and two weeks after the 6-day stimulation protocol, we re-evaluated grooming. The total time the animals spent self-grooming was scored from video recordings using ANY-maze software (version 6.0; Stoelting, Wood Dale, IL). To assess locomotor effects, the total distance traveled during the 5-min opto-stimulation (stim) for the 6-day stimulation protocol was scored automatically from video recordings using ANY-maze.

### Marble burying test

The experimental protocol for the marble burying test was based on that used previously (24). The apparatus used in this experiment consisted of a polypropylene cage (40 × 28 × 20 cm) with the floor filled approximately 5 cm deep with bedding, lightly tamped down to make a flat, even surface. A regular pattern of 20 1.5 cm diameter translucent light-blue glass marbles was placed on the surface, evenly spaced. Each animal was placed individually in the center of the cage and left for 15 min. At the end of the test, the rat was removed, and the number of marbles buried with bedding (to at least two-thirds their depth) during this interval was counted by three raters who were blind to the treatment group assignment.

### Nestlet shredding test

The experimental protocol for the nestlet shredding test was based on that described by Angoa-Pérez et al. (39). The apparatus used in this experiment consisted of a polypropylene cage with the floor filled approximately 5 cm deep with bedding and containing a cotton square of known weight that was placed on top of the bedding. The animal was placed individually in the center of the cage and left for 15 min. At the end of the test, the rat was removed, and the largest remaining intact portion of the cotton square was removed and allowed to dry. After 48 h, this portion of the cotton was weighed to calculate the percentage of nestlet shredded.

### Operant attentional set-shifting task

All set-shifting sessions were performed in standard automated operant chambers (25 × 29 × 25 cm) with metal sidewalls, a transparent Plexiglas rear wall and front door, and stainless-steel rod floor (Coulbourn Instruments, Holliston, MA), enclosed inside sound-attenuating boxes (Med Associates, Chicago, IL). A 10-W light bulb provided light throughout the sessions. Each behavioral chamber was equipped with three nosepoke holes at one of the sides of the chamber, housing infrared sensors to detect head entries and white LEDs to provide visual cues. A pellet dispenser that delivers food reward (45 mg grain-based pellets) was placed on the opposite wall. Software and an appropriate interface (GraphicState 4.0, Coulbourn Instruments) controlled the presentation and sequencing of stimuli. Behavior was recorded by video cameras mounted on the top of each unit.

The set-shifting task requires the rats to shift their response patterns between two distinct perceptual discrimination rules, or dimensions: light or side (right or left). On all trials, one of the two peripheral nosepoke holes was illuminated. Performance according to the light discrimination rule required the rats to poke the illuminated nosepoke hole, regardless of its spatial location. Performance according to the side rule required that the animals respond only at the nosepoke hole at a designated spatial location (either the left or the right) across trials, regardless of which one was illuminated (Fig. 1C).

The set-shifting protocol was modified from Darrah et al. (59). Rats were submitted to a shaping period of 5 days before training and test trials. On the first day of shaping, they were placed in the operant chambers, 10 reward pellets were delivered in the food tray at the beginning of the session and the animals remained in the chamber for 20 min. On the following day, they remained in the chamber for 20 min and during this period, reward pellets were delivered at 30-s intervals, if the rat had consumed the prior pellet. On days 3-5, rats were submitted to a single session in one of the two discrimination rules to be used in the set-shifting task, light or side (right or left). During these dimension shaping sessions, the rats were reinforced with a single reward pellet for each correct response according to the current discrimination rule. The sessions lasted until a performance criterion of 10 consecutive correct responses was reached. After shaping, in which rats learned the light and side rules individually, rats received 4 days of training sessions on the full set-shifting task. Each trial began with illumination of the central nosepoke hole; poking this central hole caused one of the two side nosepoke holes to illuminate. A correct response according to the current rule (side or light) generated a food reward; incorrect responses were not rewarded. Rewarded and non-rewarded responses were followed by a 7-s interval and the beginning of a subsequent trial. When a rat reached the performance criterion of 5 consecutive correct choices in the first rule, the rewarded dimension shifted immediately to the alternative rule (extradimensional shift), requiring the rat to shift its behavior to receive a reward. No additional cue, besides the absence of reward after a wrong response, indicated the change in rules. During each training day, the task required the rats to reach the performance criterion eight consecutive times, resulting in seven consecutive extradimensional shifts. After training, animals were submitted to six baseline testing sessions to establish a baseline performance level for each individual rat. The test session followed the same protocol as the training sessions. After the baseline testing phase, the rats were removed from the food deprivation regimen and, when the original weight was reached, approximately within a week, animals were submitted to the optostimulation. After the 6-day opto-stimulation protocol, rats were again food restricted and, after reaching target weight, underwent set-shift testing during 6 consecutive days with the same parameters as those described for the baseline testing.

### Measuring OFC-VMS connectivity via evoked potentials

Repeated optogenetic stimulation of OFC terminals in VMS is believed to induce synaptic plasticity in mice. To determine whether this effect holds in rats, we evaluated the bidirectional OFC-VMS evoked response potential (ERP) before and after the 6-day optostimulation protocol (Fig. 1D). For this, animals with active virus in the OFC-VMS pathway and electrodes in both regions (Fig. 1E) were submitted to the same optogenetic stimulation protocol previously described. Before running this experiment, rats were connected to the patch cable and stimulation/recording wires for 4 days while in the test chamber (15 min per day) for habituation, without receiving stimulation. To measure OFC-VMS effective connectivity, a PC running a custom-made LabVIEW program (National Instruments, Austin, TX) was connected to a NI USB-6343 BNC analog/digital interface unit (National Instruments), and an analog stimulus isolator (model 2200, A-M Systems, Sequim, WA). The software delivered constant-current, single pulses through the stimulus isolator. From the fifth to the eleventh day, after animals were connected to patch cable and wires, 30-50 pulses (200 uA, 0.1 ms, with an interpulse interval varying between 3 and 4 s) were delivered to OFC or VMS. The distant response to these pulses (in VMS to OFC stimulation, or in OFC to VMS stimulation) is a measure of synaptic strength between the two regions. Next, the same optogenetic stimulation protocol followed (5-min pre-stim grooming observation, 5-min stim, 5-min post-stim grooming observation; 10 ms pulses, 10 Hz, 7 mW). After the post-stimulation period, a second measurement sequence of 30-50 pulses to each area was administered. During the whole session, LFP signals were recorded continuously at 30 kHz with an Open Ephys acquisition board (60) and an Intan 32-channel headstage (RHD2132, Intan Technologies, Los Angeles, CA). Broadband activity was recorded, stored for offline analysis, filtered (band pass between 0 and 20 Hz by the amplifier), and post-processed in Matlab (Mathworks, Natick, MA). To measure inter-regional ERPs, we calculated the absolute value of the area underneath the curve (AUC) after each pulse, beginning at the offset of the stimulation artifact and measured out to 1.5 s post-stimulation.

### Histology and Immunostaining

At the end of the experiments, optical stimulation was performed for 5 min to induce immediate-early gene activation consistent with that induced by the main intervention. Ninety minutes later, rats were deeply anesthetized (pre-anesthesia with isoflurane followed by Beuthanasia-D Special, 150 mg/kg), and then transcardially perfused with cold phosphate buffered saline (PBS) followed by 4% paraformaldehyde (PFA) in 0.1 M PBS solution. Brains were extracted and kept in PFA for 24-48 h, transferred to 30% sucrose PBS for 48-72 h, flash frozen in −75 °C isopentane, and then 40 μm sections were obtained using a cryostat. To verify fiber placement, sections were Nissl-stained. Expression of ChR2-eYFP or eYFP was confirmed using fluorescence microscopy (Keyence, Osaka, Japan). Five rats showing no eYFP expression in OFC and/or showing fiber placements outside VMS were excluded from the analysis (one rat from the control group and four rats from the ChR2 group).

For c-Fos immunostaining, sections were washed three times in PBST (PBS with 0.2% Triton-X) and incubated in PBST+NGS (5% normal goat serum) for 1 h. Rabbit anti-c-Fos (1:5000; part number 5348, Cell Signaling Technology, Danvers, MA) was incubated overnight at 4 °C. The next day, sections were washed 3 times in PBST, followed by 2-h room temperature incubation in Alexa 647 donkey anti-rabbit antibody (part number 711-606-152, Jackson ImmunoResearch, West Grove, PA). Sections were then washed and processed using 4”,6-diamidino-2-phenylindole (DAPI, 1:10000) for 20 min, followed by another PBS wash. Sections were mounted on slides, using VectaShield mounting medium, and coverslipped. Histological counting of cells expressing c-Fos protein in a 720 × 540 μm area of regions of interest was performed using FIJI (61).

### Analysis of results

Statistical analysis was performed using R (version 3.6.0, R Core Team, 2019) with the package lme4 (version 1.1-23; 62). For each test, generalized linear mixed models (GLMMs) were used to account for the repeated measures design and non-Gaussian distribution of the data. For the countable data (marble burying test, number of errors in the set-shifting task, positive c-Fos cells) GLMMs with log link function and Poisson distribution were used; for the other measurements, GLMMs with identity link function (locomotion, nestlet shredding test, reaction time in the set-shifting task, ERP measurements) or log link function (grooming) and Gamma distribution were used. For each model, we used the rat”s identity as a random factor and Treatment (Control and ChR2) and/or Phase (e,g, pre- and post-stimulation) as fixed factors.

## Results

### Virus and fiber location

Viral expression was largely restricted to the medial and ventral OFC (Fig. 2A). Tracks of the optical fibers were identified in the VMS (Fig. 2B). ChR2-eYFP expression was confirmed in OFC (Fig. 2C) and in OFC projections in VMS (Fig. 2D), as targeted by Ahmari et al. (20).

**Fig. 2.**
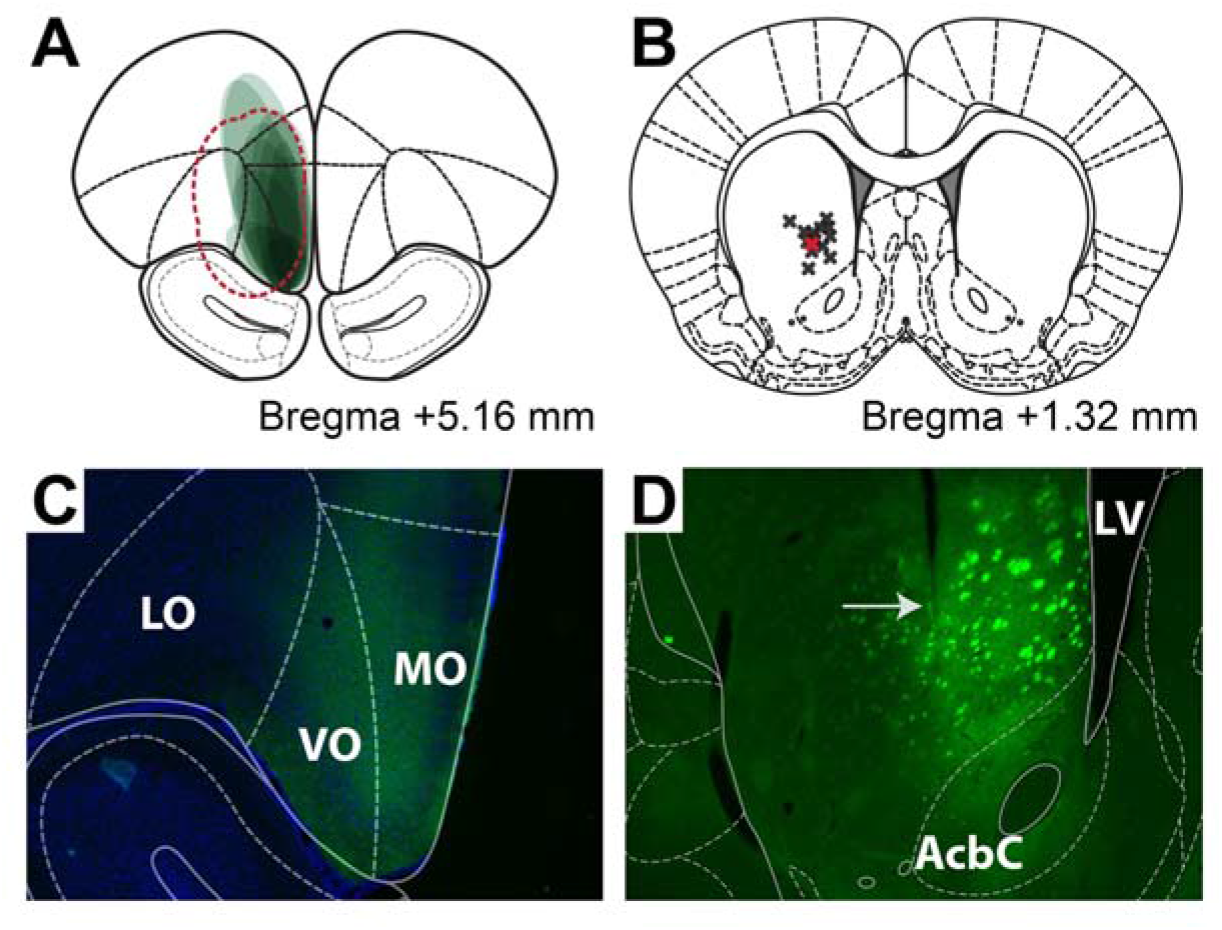
OFC-VMS projection targeting. **A)** Schematic representation of eYFP expression in the orbitofrontal cortex (OFC). Each ellipse represents the expression region of a single animal. Red dashed line depicts the virus spread in the original study (20). **B)** Localization of the optical fibers in the ventromedial striatum (VMS). Red cross represents the optical fiber”s approximate location in the original study (20). **C)** Example DAPI stained coronal section showing eYFP expression (green) in the OFC. LO = Lateral orbitofrontal cortex; VO = Ventral orbitofrontal cortex; MO = Medial orbitofrontal cortex. **D)** eYFP expression in the VMS projections directly below the optical fiber track (arrow), after virus injection into the OFC. AcbC = Accumbens nucleus core; LV = Lateral ventricle. Modified from Paxinos and Watson (63).

### Self-grooming behavior

In contrast to the prior mouse results (20), OFC-VMS repeated stimulation did not increase self-grooming in rats. There was no significant effect of active vs. eYFP-only virus on self-grooming, either before (p = 0.68), during (p = 0.80), or immediately after the stimulation (p = 0.85; Fig. 3A). Stimulation did not affect locomotor behavior (p = 1.00; Fig. 3B). There was no significant difference between ChR2 and control groups on selfgrooming evaluated one hour after stimulation (p = 0.80; Fig. 3C). We similarly did not replicate the finding that stimulation leads to long-term grooming changes; there were no differences on grooming expression one or two weeks after the end of the optogenetic stimulation protocol (p = 0.11; Fig. 3D).

**Fig. 3.**
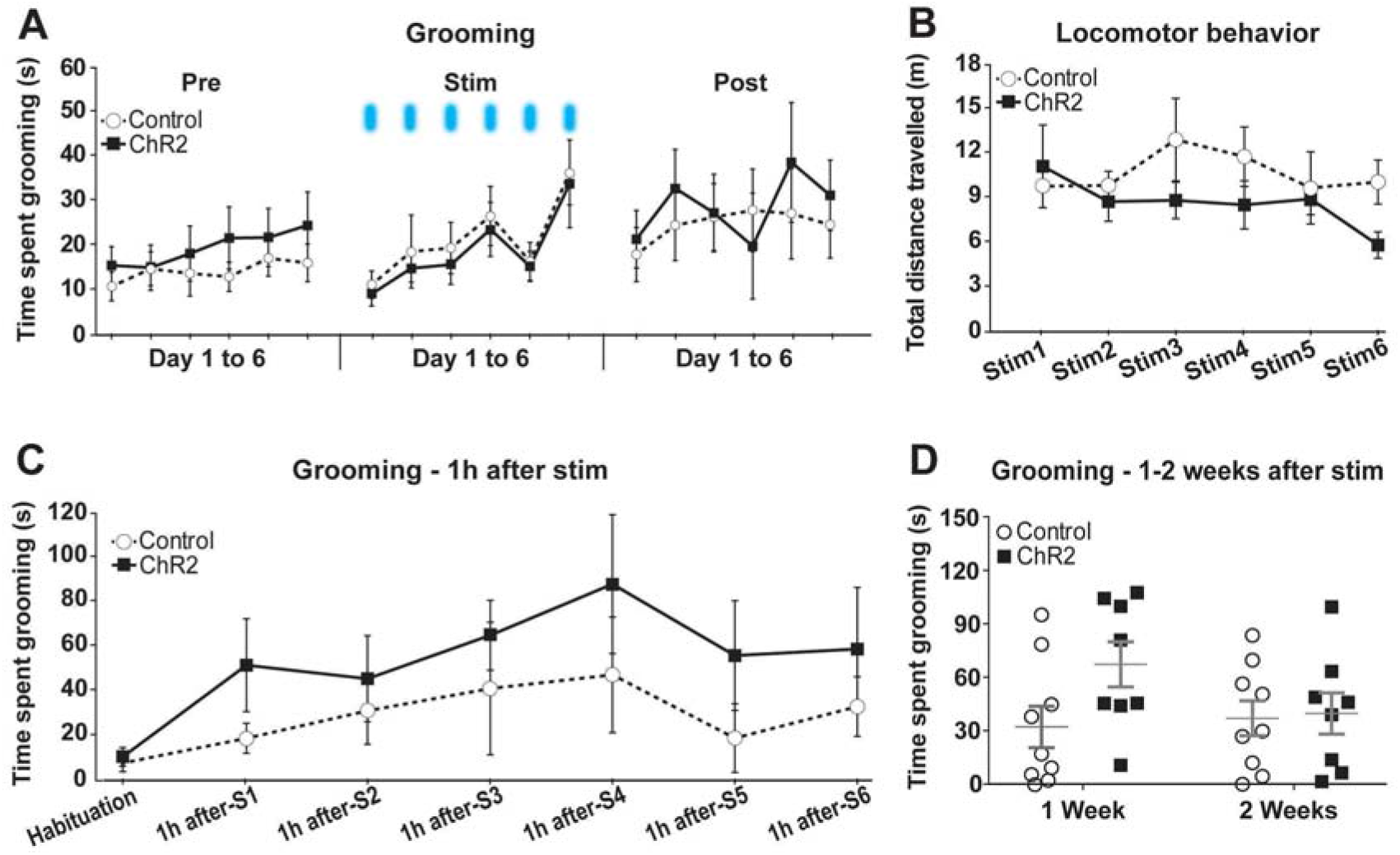
OFC-VMS repeated opto-stimulation did not increase self-grooming or locomotor behavior in rats. **A)** Total duration of self-grooming behavior assessed for 5 min immediately before (Pre), during (Stim), and immediately after (Post) 6 days of OFC-VMS opto-stimulation. **B)** Total distance traveled during the 5-min opto-stimulation over six consecutive days. **C)** Total duration of self-grooming behavior, assessed for 5 min at one hour after OFC-VMS optostimulation, for six consecutive days. **D)** Total duration of self-grooming behavior, each assessed for 5 min, one and two weeks after the end of the 6-day stimulation protocol. Markers in panels A-C represent mean, error bars show SEM. In panel D, markers represent individual animals, bars indicate mean and SEM. n = 9 for ChR2 and n = 8 for Control.

### Marble burying and nestlet shredding

OFC-VMS stimulation did not increase marble burying or nestlet shredding. There was no effect of active vs. eYFP-only virus on the marble burying test (p = 0.42; Fig. 4A). Marble burying, in fact, decreased over time in both the active stimulation and the control groups 1 h (exp(β) = 0.36, SE = 0.22, z = −4.60, p = 4.21e-6), 1 week (exp(β) = 0.48, SE = 0.20, z = −3.66, p = 2.54e-4, and 2 weeks (exp(β) = 0.07, SE = 0.46, z = −5.93, p = 3.11e-9) after stimulation in comparison to baseline. There was no effect of active vs. eYFP-only virus on the nestlet shredding test (p = 0.22; Fig. 4B).

**Fig. 4.**
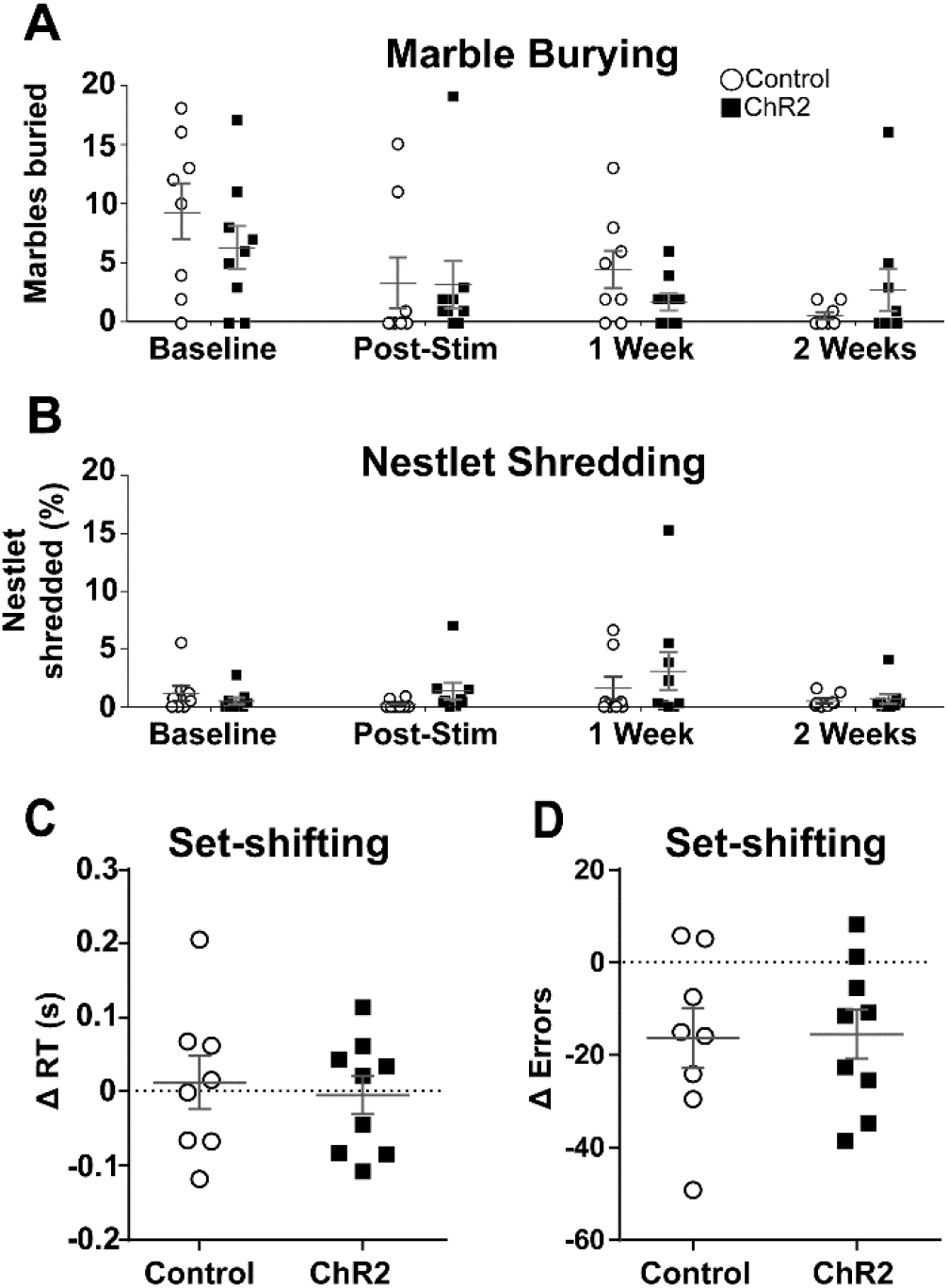
No effect of OFC-VMS opto-stimulation on non-grooming compulsive behaviors. **A)** Number of marbles buried before (Baseline) and after (Post-Stim, 1 and 2 Weeks) the 6-day stimulation protocol. **B)** Percentage of nestlet shredded before (Baseline) and after (Post-Stim, 1 and 2 Weeks) the stimulation protocol. **C)** Change in reaction time (RT) during the set-shifting sessions, from before to after the 6-day stimulation protocol. **D)** Change in the number of errors during the set-shifting sessions from before to after the stimulation protocol. Markers represent individual animals; bars indicate mean and SEM. n = 9 for ChR2 and n = 8 for Control.

### Operant attentional set-shifting task

OFC-VMS stimulation did not affect set-shifting performance. There were no active vs. eYFP-only virus differences in reaction time (p = 0.08; Fig. 4C) or number of errors (p = 0.60; Fig. 4D) before and after the 6-day stimulation protocol.

### Measures of circuit engagement

The preceding results could be explained if this stimulation protocol was insufficient to engage and activate OFC-VMS circuitry. We verified, however, that the circuitry was activated, and that OFC-VMS repeated stimulation did change their functional interaction in rats. There was an increased c-Fos expression in the OFC after opto-stimulation of the VMS in ChR2 animals compared to eYFP-only controls (exp(β) = 1.8, SE = 0.20, z = 2.88, p = 0.004; Fig. 5A-B). There was no difference for c-Fos expression in the VMS (p = 0.56; Fig. 5C).

**Fig. 5.**
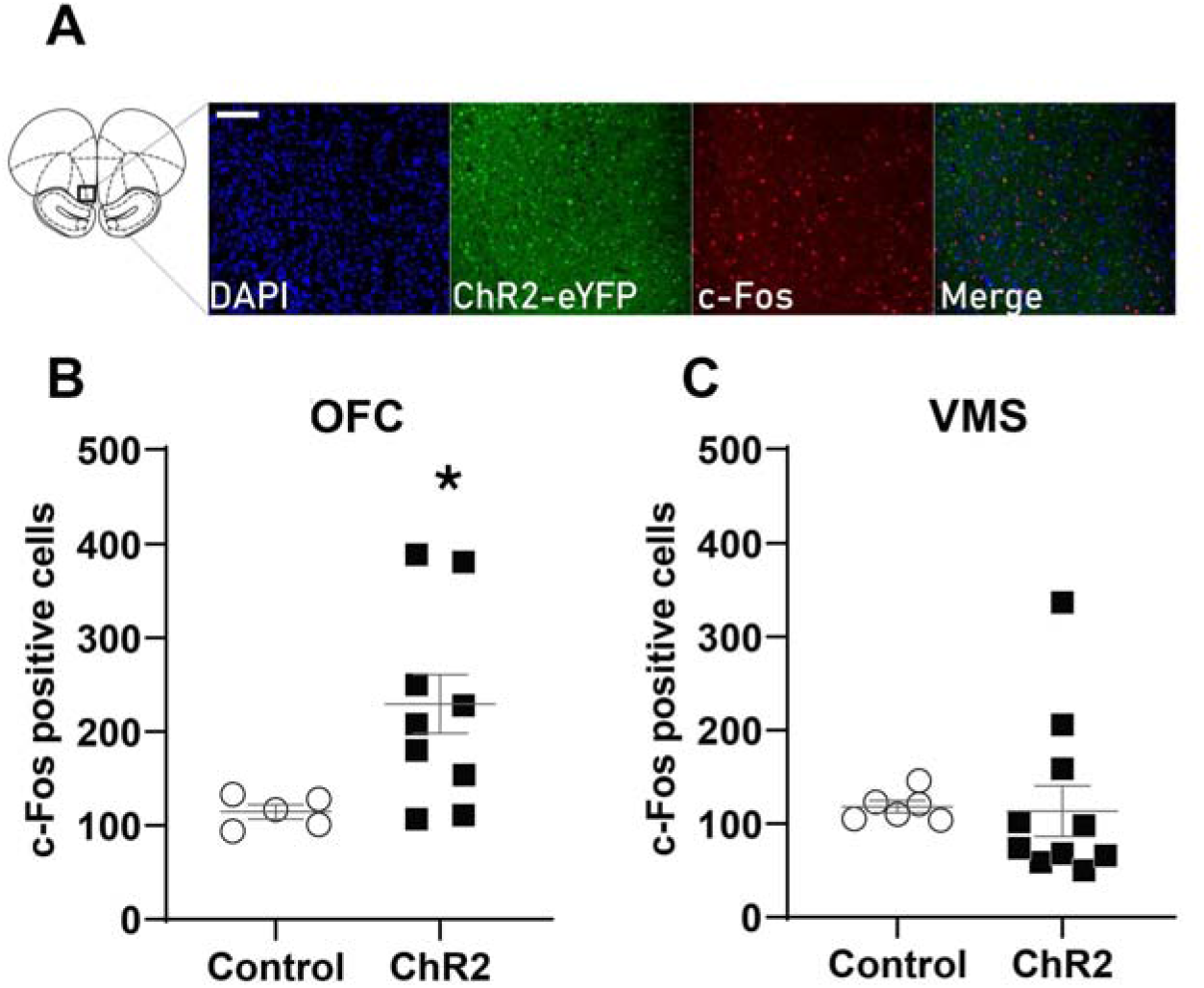
VMS opto-stimulation increased c-Fos expression in the OFC. **A)** Representative images of a slice of the OFC demonstrating light-induced neural activation of OFC through optical stimulation of the VMS. DAPI labeling (blue), ChR2-eYFP viral expression (green), and c-Fos immunostaining (red). Scale bar 100 μm. **B)** Quantification of c-Fos-positive cells in the OFC of ChR2 animals compared to eYFP-only controls. **C)** Quantification of c-Fos-positive cells in the VMS of ChR2 animals compared to eYFP-only controls. Markers represent individual animals; bars indicate mean and SEM. OFC: n = 9 for ChR2 and n = 5 for Control. VMS: n = 10 for ChR2 and n = 6 for Control. Modified from Paxinos and Watson (63).

We ran a second experiment in which we verified this activation electrophysiologically. In ChR2-transfected animals, there were clear electrical responses to light pulses (Fig. 6A-B). We were also able to measure ERPs at OFC and VMS in response to an electrical pulse delivered to the other structure, verifying *in vivo* connectivity (Fig. 6C-D). For the OFC electrical stimulation and VMS recording (Fig. 6E), when compared to Day 1, the responses increased significantly at Day 2 (β = 0.85, SE = 0.21, z = 4.01, p = 5.97e-5), Day 3 (β = 0.63, SE = 0.22, z = 2.89, p = 0.004), Day 4 (β = 1.49, SE = 0.28, z = 5.26, p = 1.41e-7) and Day 5 (β = 1.16, SE = 0.23, z = 5.15, p = 2.61e-7) of optical stimulation. Similar to Ahmari et al. (20), this plasticity was most evident across days – there was only a significant intra-day change from pre to post stimulation on testing Day 4 (β = −0.75, SE = 0.36, z = −2.08, p = 0.04). For the VMS electrical stimulation and OFC recording (Fig. 6F), there were changes only at Day 3 of optical stimulation in comparison to Day 1 (β = 2.11, SE = 0.71, z = 2.96, p = 0.003). This second cohort again had no significant change in self-grooming behavior from OFC-VMS repeated stimulation (p = 0.76; Fig. 6G).

**Fig. 6.**
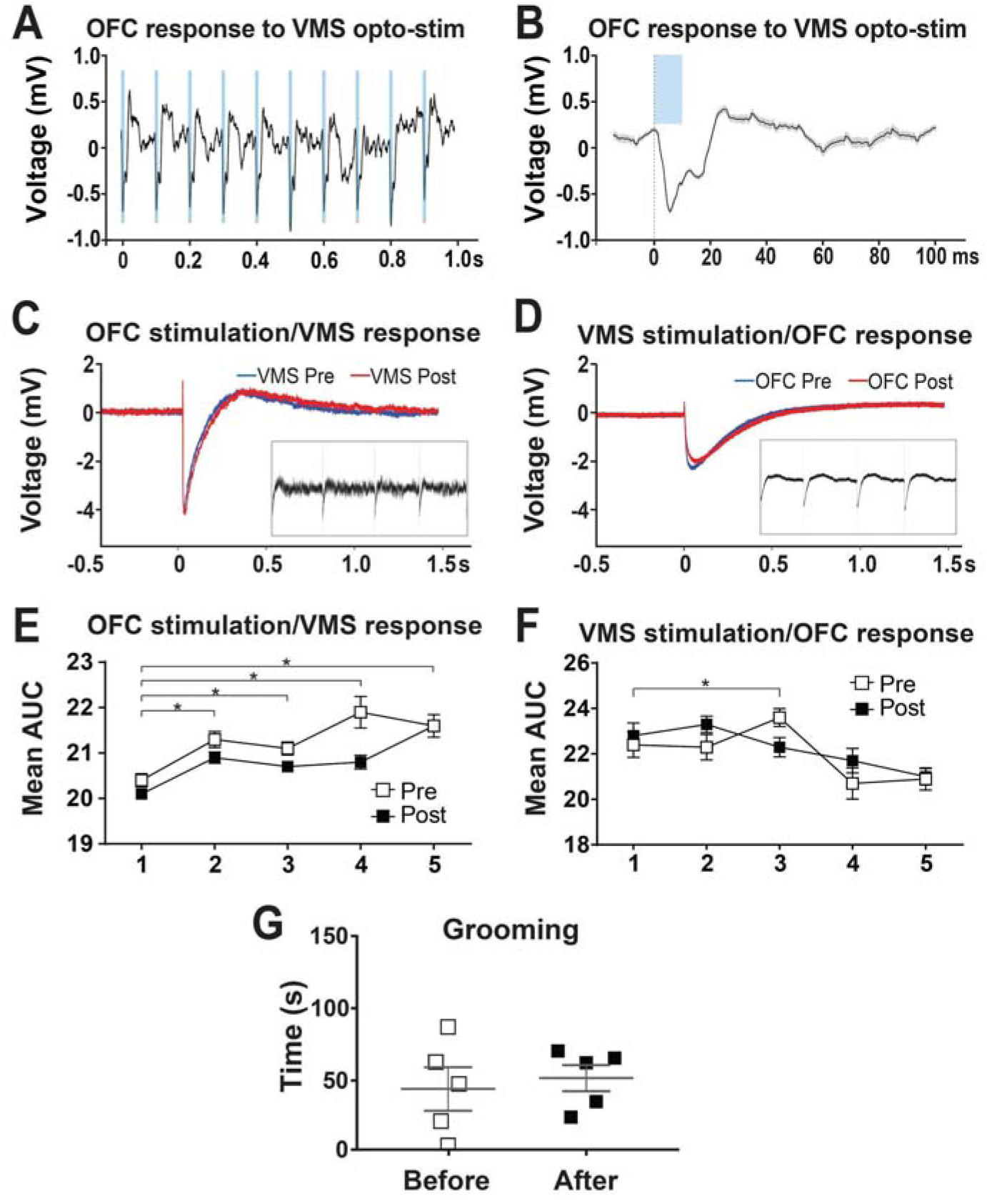
OFC-VMS repeated opto-stimulation induced plasticity in rats. **A)** In vivo recording in awake rats shows field responses in the OFC to optical stimulation of VMS projections. **B)** Singletrial example of the population response shown in (A). **C)** Representative evoked response in VMS to OFC electrical pulses, pre- and post-repeated opto-stimulation. (Inset) In vivo recording in awake rats shows field responses in the VMS to electrical stimulation of OFC. **D)** Representative evoked response in OFC to VMS electrical pulses, pre- and post-repeated opto-stimulation. (Inset) In vivo recording in awake rats shows field responses in the OFC to electrical stimulation of VMS. **E)** VMS response, over time, to OFC stimulation, showing increase in this connectivity metric between days, but not within each day. **F)** OFC response, over time, to VMS stimulation, showing increase in connectivity only for day 3 of opto-stimulation. Note that E-F plot the area under the ERP curve calculated from individual stimulation trials across all rats, while C-D plot mean ERPs from single animals. As such, group-level differences are not readily visible in C-D. **G)** OFC-VMS repeated opto-stimulation did not increase self-grooming in this second cohort of rats. Markers in panels E and F represent mean, error bars show SEM. In panel G, markers represent individual animals, bars indicate mean and SEM. n = 5.

## Discussion

We could not reproduce in rats a model of compulsive behavior that has been previously reported in mice (20), suggesting that the OFC-VMS pathway is not sufficient for compulsivity. In a cohort that was explicitly and strongly powered to detect effect sizes below those of the original report, repeated OFC-VMS opto-stimulation did not increase the expression of self-grooming behavior in rats. This is inconsistent with a previous report for mice, in which the cortico-striatal stimulation led to a significant increase in selfgrooming immediately before and one hour after stimulation (20). We also extended the evaluation of the potential effects of the OFC-VMS stimulation to other common assays of compulsivity. The repeated OFC-VMS stimulation protocol did not affect marble burying, nestlet shredding, or operant set-shifting behaviors in rats either. In mice, acute OFC-VMS stimulation led to a large but transient increase in locomotion compared with controls, followed by a return to baseline levels after stimulation (20). In rats, there was no significant difference for the animals” locomotor behavior during the 5 min of stimulation over the six days of our protocol.

Critically, we saw no behavior change even though the protocol was successful in remodeling the OFC-VMS circuitry. Optogenetic stimulation of the OFC-VMS pathway caused acute neuronal activation, as measured by electrical responses to light pulses and by immediate early gene expression. Further, we reproduced the directional synaptic plasticity of the original mouse report: stimulation was also sufficient to induce changes in synaptic weighting in the OFC-VMS direction, but caused minimal changes in the VMS-OFC direction. This implies that in rats, and presumably in higher species, compulsive behavior cannot be attributed to simple monosynaptic changes. Although not directly tested in this preparation, the same may be true for other monosynaptic findings reported in mice, e.g., the role of similar cortico-striatal circuitry in addiction (64).

This translational failure emphasizes the need to carefully consider species differences and suggest cautiousness about overgeneralizing phenomena observed in a single species to human illness. Rodents are a vital model system for neurobiology due to the ability to use more invasive and comprehensive techniques (65–68). Experiments that require altered protein expression in specific neurons most often use mouse models, due to the greater flexibility of genetic targeting. Physiological, anatomical and psychopharmacological studies, however, have predominantly been performed in rats (69–71). Although closely related, mice and rats present important genetic, anatomical, physiological, and behavioral differences that complicate direct comparisons between species (68,72). The differences between our results and those of Ahmari et al. (20) may in part be explained by environmental factors such as housing enrichment, learning, and stress that can all affect behavior. Given that we saw comparable physiologic changes, however, we believe it is more valid to conclude that findings in the CSTC circuitry of mice may not necessarily be applicable in rats, and thus may not translate to species higher on the phylogenetic chain.

These results do not appear to be attributable to minor variation in optogenetic technique, given the parallel physiologic changes across species. We used the same viral serotype as in the original study. Variations in transduction could then be linked primarily to the promoter or to potential differences in brain circuit complexity between mouse and rat. We used a different promoter (CaMKIIa, compared to the elongation factor-1 alpha, EF1α, used in Cre mouse lines in the original study), so the levels of expression could differ across the two animal models. However, given the less common use of Cre-driver lines in rats, several studies have shown that the CaMKIIa promoter can successfully engage excitatory neurons in cortical and other brain regions (73–78). Our results cannot be explained by differences in viral delivery, since in our study a higher volume (1 μL) at a higher genomic titer (4.1×10^12 vs 1×10^12 vg/ml) was administered to account for the larger brain of rats. Fluorescence imaging of ChR2/eYFP immunostaining at VMS shows that the region targeted by the optogenetic stimulation received transfected fibers originating at the OFC (as seen in Fig. 2). Our results similarly cannot be explained based on the light delivered to the VMS. Compared to the original study, we used higher light intensities to try to mimic the behavioral response. Ahmari et al. (20) used a solid-state 473 nm 100 mW laser, with light intensity set to 1-5 mW illumination at the fiber tip in the brain. In our approach we used a 465 nm blue LED capable of generating 7 mW at the tip of the fiber, which corresponds to 55 mW/mm^2^. Light power densities of 2-5 mW/mm^2^ with a wavelength from 465-475 nm are sufficient to stimulate action potentials in neurons expressing ChR2 at typical experimental levels (80). Finally, related to both these points, there was a clear difference in neural excitation between ChR2 and eYFP-only rats. In addition to the c-Fos results, our electrophysiological study confirmed that optogenetic stimulation of OFC-transfected projections in VMS acutely affected neuronal physiology (Fig. 6).

This failure of translation between rodent species suggests a need for caution when attempting to model compulsivity or other CSTC-linked human phenomena. There is functional homology across rodents and primates in at least three different regions of the striatum: the dorsal regions being specialized in motor aspects, such as planning and execution; the medial region involved in learning and attention; and the ventral region having a prominent role in reward and emotion processing (12,80–83). Dopaminergic system organization in the striatum is also conserved across rats and mice (84). However, the prolonged development and more complex and flexible behavioral repertoire of the rat compared with mice (85) suggest that anatomic homology may not fully reflect circuit function. For example, rats use more complex strategies than mice do to locate the platform in the Morris water maze and, consequently, perform better than mice in learning spatial information (85–87). Inhibition of adult neurogenesis in the hippocampus produces deficits in contextual fear conditioning behavior in rats but not mice (88). Additional knowledge of functional homology, e.g. at the physiologic level, might be necessary for more accurate translational studies (84). It may also be necessary to consider the role of timing-dependent plasticity. Neither our work nor Ahmari et al. (20) captured the timing of optogenetic stimulation relative to spontaneous grooming bouts. It may be that, by pure chance, the mice in the original study groomed frequently enough that optogenetic stimulation tended to coincide with their grooming. This might have caused positive, self-reinforcing associations that led grooming to become habitized and over-expressed. With the advent of high-precision automatic behavior estimation tools (89,90), this hypothesis could be tested in future work.

Taken at face value, our findings fit into an ongoing evolution in circuit theories of compulsivity. Initial CSTC hyperconnectivity theories emphasized OFC-originating circuits, including pathways targeting the ventral striatum (19,91,92). There was a strong emphasis on anxiety and other negative-valence affective constructs. Those ideas are now evolving, with an increasing emphasis on more dorsal CSTC circuits involved in executive function and planning (17) and on the role of other structures such as amygdala (92). Part of that evolution is a growing understanding that CSTC circuits are not segregated, but implement a spiral-like transfer of information between recurrent cortico-basal loops (17,19,93). Although not yet proven, this interconnection between loops may increase with increasing brain size/complexity. As a result, grossly observable behaviors might be elicited in mice if the OFC-VMS circuit acts in relative isolation, but greater cross-loop integration in rats and primates might buffer or compensate for a circuit-limited perturbation, and the same OFC-VMS circuit change might no longer produce behavior change. If true, this model suggests that for clinically severe compulsivity to emerge in a human brain, the CSTC circuitry might require multiple simultaneous “hits” that remove the capacity for compensation. Thus, a more translatable model might study the effect of multiple circuit disruptions in conjunction, or might combine this type of single-synapse manipulation on a background of either genetic loading or early life adversity. In that sense, our failure to translate this specific protocol highlights the need for rodent models that more fully capture the human pathway to illness.

## Acknowledgements

Research has been supported by the São Paulo Research Foundation (FAPESP, Brazil - 2017/22473-9); OneMind Institute; MnDRIVE Brain Conditions Initiative; and UMN Medical Discovery Team on Addictions.

